# Experimental and methodological framework for the assessment of nucleic acids in airborne microorganisms

**DOI:** 10.1101/2023.10.10.561683

**Authors:** Raphaëlle Péguilhan, Florent Rossi, Olivier Rué, Muriel Joly, Pierre Amato

## Abstract

Studying airborne microorganisms is highly challenging due to ultra-low and spread biomass, and great spatial and temporal variabilities at short scales. Aeromicrobiology is still an emerging discipline of environmental microbiology, and some of the basic practices (replication, control of contaminants, *etc*) are not yet widely adopted, which potentially limits conclusions. Here we aim at evaluating the benefits of such practices in the study of the aeromicrobiome using molecular-based approaches, and recommend the following: (i) sample at high airflow rate, if possible into a fixative agent, in order to be able to capture specific situations ; (ii) replicate sampling and process samples individually to enable statistical analyses ; (iii) check for contaminants at different steps of the analytical process, and account for their potential stochasticity in sequence decontamination methods ; (iv) include internal references to verify qualitative and quantitative aspects of the data, and (v) eventually investigate multiple analytical procedures to identify potential impacts on the data. In our study, samples were collected at a remote mountain site using high-flow rate impingers collecting airborne material into nucleic acid preservation buffer. As high of ∼75% of the sequences were shared between independent triplicates, gathering 28 to 38% of the richness observed at the ASV level at a given sampling date, which also emphasizes spatial heterogeneity at short scale due to rare taxa. Thanks to replicates, daily variations in the diversity of bacteria could be distinguished statistically, and the inevitable presence of contaminating sequences in controls could be accounted for using established statistical methods. This work opens new perspectives and notably paves the way to untargeted molecular methods in the exploration of aeromicrobiome’s composition and functioning.

**Graphical abstract:** 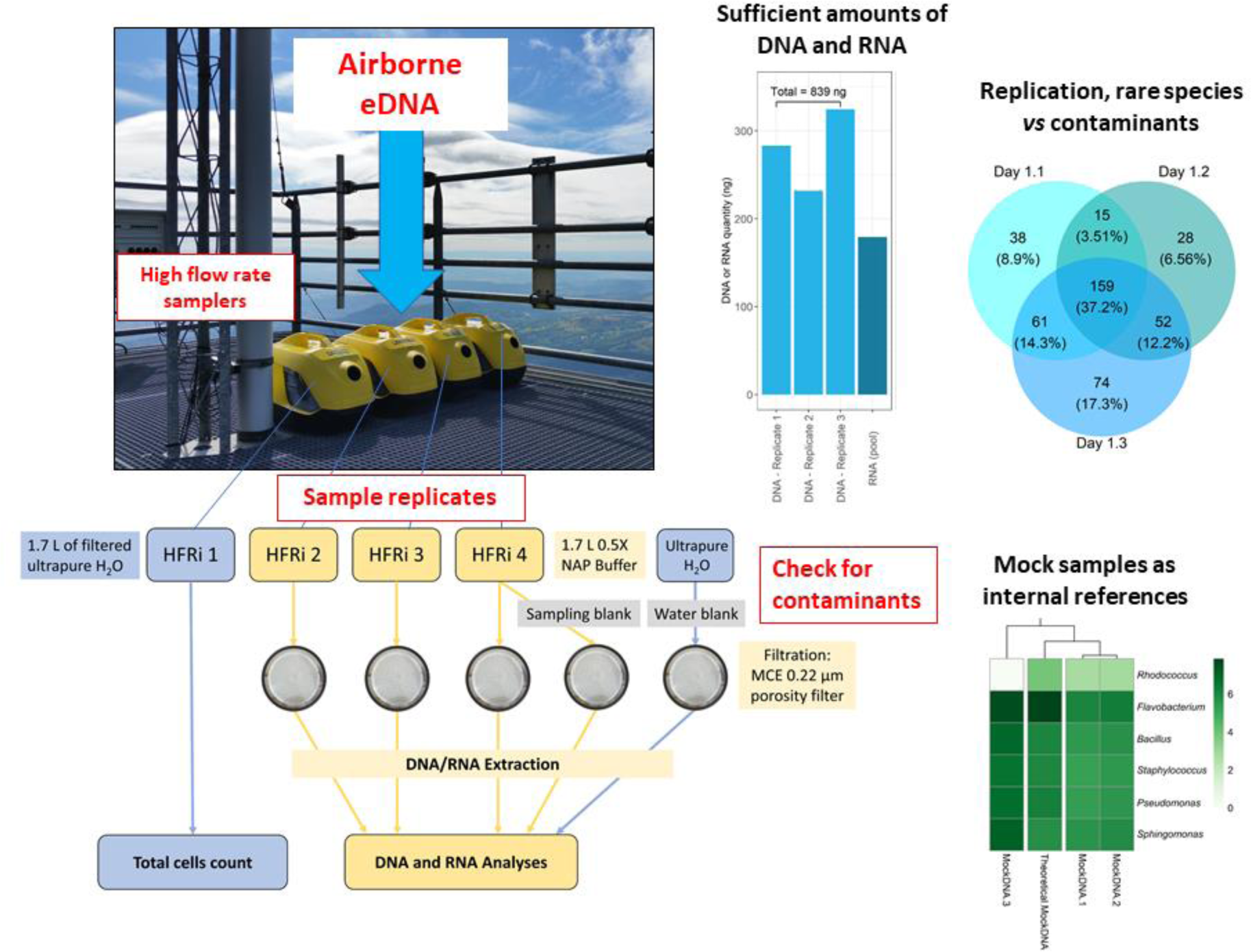

## Introduction

The atmosphere is exposed to emissions of biological material from surface ecosystems and carries their DNA imprints. These can be lifted up to high altitudes and travel over long distances up to continental scales (Després et al., 2012; Smith et al., 2013; Barberán et al., 2015). Biological aerosols play roles as cloud condensation nuclei (CCN) and ice nuclei (IN), and can have influence on cloud microphysics and precipitation (Zhang et al., 2021; Bauer et al., 2003; Möhler et al., 2007; Patade et al., 2021). Bioaerosols also include microorganisms of sanitary concern, such as pathogenic, opportunistic, or antibiotic resistant taxa (Rossi et al., 2022; Fernstrom et al., 2013; Beattie and Lindow, 1995), and, regardless of the potential hazards they may create, the fact that microbial cells can maintain metabolic activity (Krumins et al., 2014; Amato et al., 2019; Šantl-Temkiv et al., 2017) raises questions about their physiological properties and their role in atmospheric chemical processes (Wirgot et al., 2017; Khaled et al., 2021; Joly et al., 2015).

Aeromicrobiology is still an emerging discipline, and the basics of recommended practices in environmental microbiology are often neglected. The first prerequisite for a meaningful analysis is to be able to distinguish target(s) from contaminants. These can originate from handling, equipment, or reagents including commercial kits (Stinson et al., 2019; de Goffau et al., 2018; Salter et al., 2014). Preventing contaminants involves, at the very least, working under adequate sterile conditions, and checking them by taking control samples at various stages of the analytical process. In addition, statistical comparison between diverse environmental situations requires replicates (Prosser, 2010; Ji et al., 2019), which also help detecting sporadic contaminants such as those brought by commercial reagents.

The greatest technical challenges in outdoor aeromicrobiology are related to the low biomass (typically ∼1 to 100 cells per liter of air), in particular in remote situations. This imposes sampling for extended periods of time, or at high-flow rate, and preserving the *in-situ* state of the material during and upon sampling (Burrows et al., 2009; Després et al., 2012; Šantl-Temkiv et al., 2020). Sampling over long periods of time alters integrity and viability of airborne biological material and living cells (*e.g.,* Manibusan and Mainelis, 2022). Moreover, as microbial assemblages vary widely over short temporal and spatial scales, sampling may have to be carried out over short periods of time (*e.g.*, a few hours or less) depending on the objectives, for instance for capturing a specific situation. Meanwhile, sufficient quantities of biological material must be collected to allow analyses, and ensure representativeness of the samples through statistics. High-flow rate sampling solutions are therefore methods of choice, and impingers, *i.e.,* liquid impactors, are considered as reference samplers for bioaerosols (Dybwad et al., 2014; Rule et al., 2007). In addition of preserving cell integrity due to gentler impaction than on solid surfaces, these allow the use of a variety of solutions for preserving viability or nucleic acids, including RNAs (Kathiriya et al., 2021; Šantl-Temkiv et al., 2017; Griffin et al., 2011; Camacho-Sanchez et al., 2013).

As analytical methods improve, deepest investigations are made possible, and methods based on high-throughput sequencing of amplicons are now widely used to explore bacterial and fungal diversity (Zhao et al., 2022; Tignat-Perrier et al., 2020b). With the advent of NGS techniques as highly sensitive methods for investigating microbial diversity, new challenges have emerged such as detection of trace contaminants, sequencing bias and artifacts (de Goffau et al., 2018). Here, we propose an experimental procedure to study biological aerosols through nucleic acid-based approaches in remote atmosphere, considering replication and accounting for inevitable contaminants, whatever their origin. Several high-flow-rate impingers (HFRi) filled with nucleic acid preservation (NAP) buffer as the sampling fluid were deployed in parallel for molecular investigations. Controls for the presence of contaminants were made at multiple steps of the experimental procedure, and quantitative aspects as well as sequence annotation accuracy were verified using artificial “mock” communities as internal references. Amplicon sequencing data indicated good reproducibility of the methods and allowed distinguishing bacteria diversity from consecutive days with good statistical confidence. The amounts of DNA and RNA collected opens unprecedented possibilities of direct sequencing of metagenomes and metatranscriptomes from atmospheric samples.

## Materials and methods

Drastic procedures were applied throughout the experimental process, including systematic decontamination of the material used (pipets, *etc*.) with detergents (RNAse away, ethanol 70% or diluted bleach), exposure to UVs, use of laminar flow hoods, systematic filtration of all the liquids before autoclave, *etc*.

### Sampling setup with High-Flow-Rate Impingers (HFRi)

Sampling of aerosols was carried out at the summit of puy de Dôme mountain (1,465 m a.s.l, France) using the facilities of the atmospheric station (Baray et al., 2020; Péguilhan et al., 2021), during three consecutive days in July 2020. Samples were collected for 6 to 6.5 consecutive hours, corresponding to 708-770 m^3^ of air collected by each sampler at each occasion, at ambient temperature ranging from 11 to 20°C and 48 to 61% humidity (**Table 1**). More details on the meteorological context, including backward trajectories of the corresponding air masses can be found at https://www.opgc.fr/data-center/public/metadata.

**Table 1:** Main characteristics of aerosol samples.

| Sample identifier | Sampling date (dd/mm/yyyy) | Number of replicates | Sampling duration (h) | Air temperature (°C)<sup>#</sup> | Relative humidity (%)<sup>#</sup> | Wind speed (m.s<sup>-1</sup>)<sup>#</sup> | Microbial cell concentration (cell.m<sup>-3</sup>)<sup>*</sup> |
| --- | --- | --- | --- | --- | --- | --- | --- |
| Day 1 | 07/07/2020 | 3 | 6.5 | 11.1 | 61 | 3.6 | 1.62×10<sup>3</sup> |
| Day 2 | 08/07/2020 | 3 | 6.1 | 14.2 | 53 | 3.1 | 1.72×10<sup>3</sup> |
| Day 3 | 09/07/2020 | 3 | 6.0 | 20.3 | 48 | 3.4 | 2.62×10<sup>3</sup> |

<sup>#</sup> Average over the sampling period; <sup>\*</sup>: inferred from concentration in the collection liquid and sampler's air flow rate.

The overall sampling strategy is summarized in **Figure 1**. Four HFRi were used in parallel from the platform of the roof of the atmospheric station. HFRi sampler is a commercial Kärcher DS 5600 or DS6 vacuum cleaner (Kärcher SAS, Bonneuil sur Marne, France) that can contain up to 1.7 L of collection liquid and operates at an airflow rate of 118 m^3^.h^-1^ (Šantl-Temkiv et al., 2017). One of the samplers was dedicated to cell quantification by flow cytometry and was filled with filtered (0.22µm mixed cellulose esters (MCE) membranes, 47 mm diameter, Dominique Dutscher; Bernolsheim, France) and autoclaved ultrapure water as the collection liquid. The other three HFRi were dedicated to nucleic acid analyses and were filled with 0.5X nucleic acid preservation (NAP) buffer (see below for preparation). For all samplers, the volume of liquid was checked individually every hour by weighing, and it was compensated for evaporation with filtered autoclaved ultrapure H_2_O when necessary, assuming unit density. Immediately after sampling (on site), the collection liquid of each individual sampler dedicated to nucleic acid analyses was filtered independently through 0.22 µm MCE membranes using sterile filtration units (Thermo Scientific Nalgene), in a laminar flow hood previously exposed to UV light for 15 min. The filters were then individually placed into 5 mL Type A Bead-Tubes (ref. 740799.50, Macherey-Nagel), added with 1,200 µL MWA1 lysis buffer and stored at - 80°C until nucleic acids extraction as specified in the corresponding section below.

**Figure 1:**
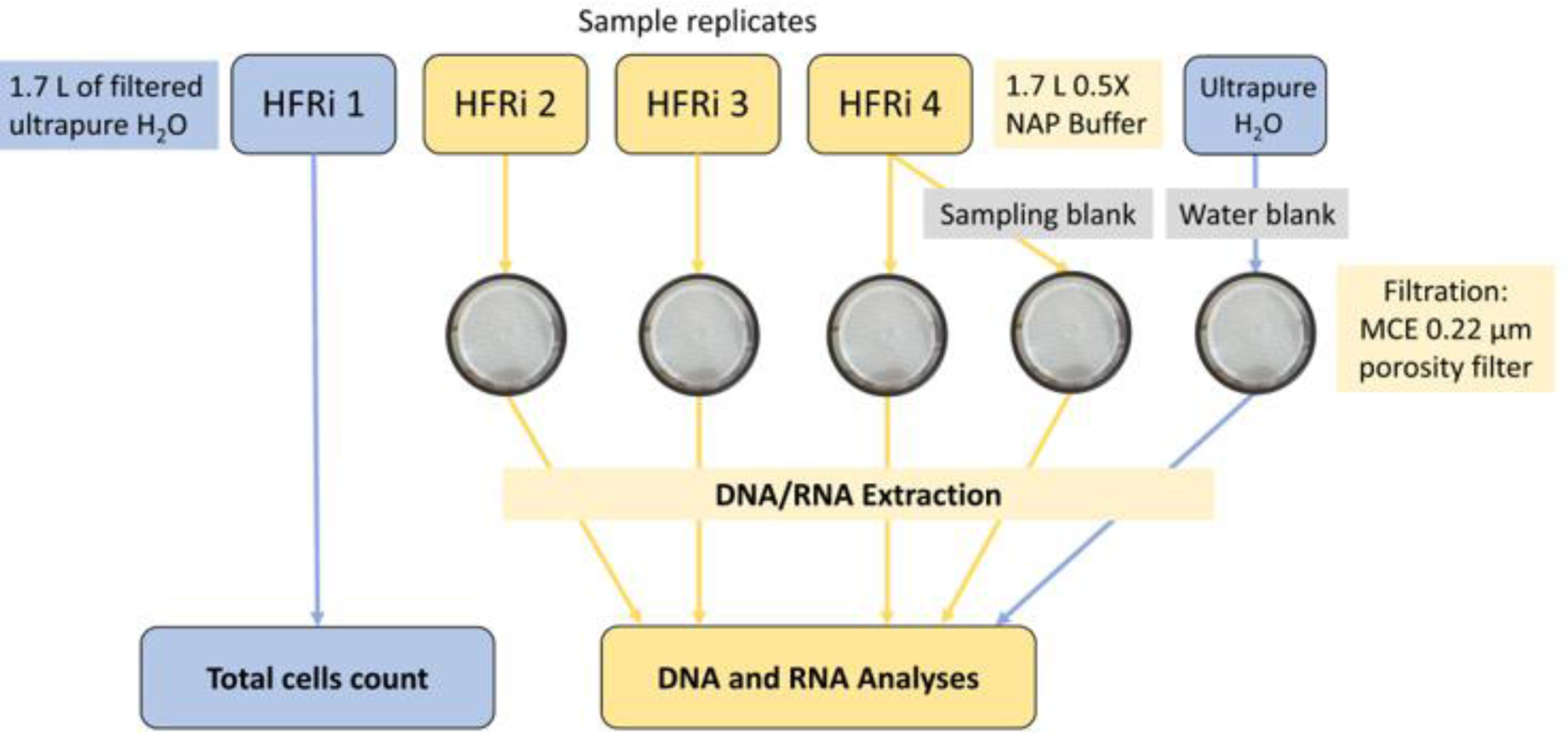
Schematic of the experimental procedure (HFRi: High Flow Rate impinger).

Negative controls of the collection liquids and samplers were performed just before sampling: 1.7 L of 0.5 X NAP buffer were poured into one of the HFRi tanks, left for 10 min with agitation, and then collected in a sterile bottle; these are referred to as “sampling blanks”. In addition, filtered autoclaved ultrapure water was used as a negative control to detect possible contamination during sample processing after collection; these are referred to as “water blanks” (**Figure 1**). All controls were processed and analyzed along with environmental samples.

NAP buffer was prepared following the instructions in Camacho-Sanchez et al. (2013). This is composed of (final concentrations): 0.019 M of ethylenediaminetetra-acetic acid (EDTA; Fisher BioReagents™), 0.018 M of trisodium citrate salt (Fisher Chemical) and 3.8 M of ammonium sulfate (ACROS ORGANICS) dissolved in ultrapure water, and H_2_SO_4_ to pH 5.2. The NAP buffer was then filtered through Glass Microfiber filters GF/F (47 mm diameter, porosity of 0.7 µm; Whatman; Maidstone, United Kingdom) to remove impurities, aliquoted and autoclaved as volumes of 1.7 L into 2-L glass bottles for subsequent direct use.

### Sampler decontamination procedure

Ethanol and UVs are common decontamination procedures (Dommergue et al., 2019; Archer et al., 2019; Šantl-Temkiv et al., 2017). Here the standard decontamination procedure of the polypropylene HFRi’s tanks consisted in: (1) thorough rinse with H_2_O then deionized H_2_O; (2) rinse and exposure to 2 L of 70% ethanol for 10 min; (3) exposure of the different parts of the tanks to UVs (254 nm) for 10 min, within a laminar flow hood. The decontaminated tanks were then stored in sterile autoclave bags until deployment to the field.

To evaluate the efficiency of our decontamination procedure, two sampling tanks were intentionally contaminated with a culture of *Pseudomonas syringae* 32b-74 (see **Supplementary Figure 1**). The strain was cultured in liquid R2A at 17°C for 3 days and then centrifuged for 3 min at 13,000 g to recover cells. The supernatant was removed and the pellet rinsed and resuspended in dH_2_O, at a concentration of ∼2.3×10^9^ cell mL^-1^ estimated from OD_600_. Finally, 7.2 mL of the stock solution was diluted in 1.7 L of autoclaved ultrapure water to reach a final solution at ∼10^7^ cell mL^-1^, and poured into sampler’s tank. Such cell concentration in the collection liquid is much higher than what can be expected from sampling in natural context (Šantl-Temkiv et al., 2017); a third sampler was left uncontaminated as a control. The samplers were then emptied and subjected to decontamination, using either the standard procedure described above, or just the thorough rinsing step with dH_2_O, then refilled with 1.7 L of filtered (0.22 µm porosity) autoclaved ultrapure water, as for actual sampling. Samplers were then switched on for 10 minutes to ensure contact with all tank’s parts, and subsamples of the collection liquid were then analyzed for total cells count as detailed below in the corresponding section. The autoclaved ultrapure water used was also analyzed without contact with the samplers. Total cell concentrations in the liquid exposed to the samplers after intentional contamination and decontamination were similar as those in uncontaminated controls (Mann-Whitney test; *p-value* < 0.05). A simple thorough rinse with filtered ultrapure water thus removed cells, and UV exposure ensured further decontamination.

### Total cell counts

Total cells were quantified by flow cytometry from triplicate subsamples of 450 µL of the collection liquid from the dedicated HFRi, fixed with 50 µL of 5% glutaraldehyde (0.5% final) and stored at 4°C until analysis, using a BD FacsCalibur instrument (Becton Dickinson, Franklin Lakes, NJ). Before analysis, samples were added with TE buffer (pH 8.0) and SYBR Green I stain following the protocols in Amato et al. (2017). Negative controls consisted of ultrapure water as the template.

### Nucleic acid extraction

Environmental samples were processed within a laminar flow hood previously exposed to UVs (15 min), and all bench surfaces, pipets etc. were decontaminated using RNase away spray solution (Thermo Scientific; Waltham, USA). Three nucleic acid extraction kits designed for DNA and RNA extraction were compared: DNeasy PowerWater kit (QIAGEN; Hilden, Germany), NucleoSpin Soil, and NucleoMag^®^ DNA/RNA Water kit for water and air sample (Macherey-Nagel, Hoerdt, France), referred to as the “Water”, “Soil” and “Air” kits, respectively. The “Water” and “Soil” kits, and the “Soil” and “Air” kits were compared as pairs during two sampling events each, in triplicate using three independent samplers. After each sampling event, the collection liquids were entirely and individually filtered through 0.22 µm MCE membranes. The filters were then cut equally into two pieces for extraction using 2 of the kits and stored at -80°C until processing. Extractions were performed following the manufacturers’ protocols. In the case of the “Air” kit, slight adaptations were made from manufacturer’s instructions: half MCE filters were placed into individual 5 mL Type A Bead-Tubes (ref. 740799.50, Macherey-Nagel) and added with 1,200 µL MWA1 lysis buffer. After the lysis step (5 min of bead-beating using a vortex), ∼600 µL of lysate was processed following the protocol adapted for 47 mm filter membranes. For DNA, the lysates were added with 1:50 volumes of RNase A (12 mg/mL, stock solution), incubated for 20 min at room temperature, then eluted into 50 µL of RNase-free H_2_O after another incubation for 5 min at 56°C. DNA in the eluates was finally quantified using Quant-iT™ PicoGreen dsDNA kit (Invitrogen; Thermo Fisher Scientific, Waltham, MA USA). For RNA, the lysates were added with rDNAse and processed as recommended. Purified RNAs were finally quantified by fluorimetry using RiboGreen (Invitrogen; Thermo Fisher Scientific, Waltham, MA USA). In terms of quantity of DNA recovered, the “Air” kit outperformed the “Soil” kit, which itself surpassed the “Water” kit (see **Supplementary Figure 2**). The “Air” kit was therefore selected for further investigations; it has the additional advantage of allowing parallel extraction of DNA and RNA.

### 16S ribosomal gene amplification by PCR

Amplification of the V4 region of the 16S rRNA gene of bacteria was performed from genomic DNA extracts by multiplexed PCR, using the primers 515f (5’-GTGYCAGCMGCCGCGGTAA-3’) (Parada et al., 2016) and 806r (5’-GGACTACNVGGGTWTCTAAT-3’) (Apprill et al., 2015). The PCR mix was composed as follows: each 50 µL reaction volume contained 2 µL of sample, 10 µL of 5X Platinum II PCR Buffer (Invitrogen; Thermo Fisher Scientific, Waltham, MA USA), 5 µL of Platinum GC Enhancer, 1 µL of 10 nM dNTPs (Sigma-Aldrich; Merck, Darmstadt Germany), 1 µl of 10 µM forward and reverse primers, 0.2 µL of Platinum II Taq HS DNA pol (Invitrogen), and 29.8 µL of Ambion™ Nuclease-free water (Invitrogen). PCR amplification conditions (35 PCR cycles) are described on the “Earth Microbiome Project” website (https://earthmicrobiome.org/). All amplicons (environmental samples and negative and positive controls) were purified using QIAquick PCR Purification kit (QIAGEN; Hilden, Germany), pooled equimolarly and sequenced on Illumina Miseq 2*250 bp (GenoScreen; Lille, France).

### Quality controls for taxonomic affiliation and biodiversity profiling

Artificial mock samples prepared from pure cultures or DNA extracts were processed as internal references, down to sequencing. An artificial atmospheric “mock” community was obtained by mixing pure cultures of six bacterial strains isolated from cloud water and representing a range of typical atmospheric bacteria (Amato et al., 2007; Vaïtilingom et al., 2012; Lallement et al., 2017): *Pseudomonas syringae* PDD-32b-74 (GenBank ID for 16S rRNA gene sequence: HQ256872), *Bacillus* sp. PDD-5b-1 (DQ512749), *Sphingomonas aerolata* PDD-32b-11 (HQ256831), *Rhodococcus enclensis* PDD-23b-28 (DOVD00000000), *Staphylococcus equorum* PDD-5b-16 (DQ512761) and *Flavobacterium* sp. PDD-57b-18 (KR922118.1). These were cultured separately in 10 mL of liquid R2A at 17°C until late exponential phase (21-44h incubation). Genomic DNA were extracted either from pure cell suspensions then mixed at known concentrations (“Mock DNA”), or from cell suspensions mixed at known concentrations before DNA extraction (“Mock Cloud”) (**Supplementary Figure 3; Supplementary Table 1**). The former allowed evaluating differences in DNA amplification efficiency depending on taxa, while the latter evaluated in particular the efficiency of DNA extraction.

For the “Mock DNA” samples, DNA extraction was performed following manufacturer’s protocol of the QIAamp DNA Mini kit (QIAGEN, Hilden, Germany) with minor changes: 1 mL of each culture was centrifuged at 14,000 g after 4 days of incubation and the pellets were re-suspended in 180 µL of TE (1X), with 25 µL of lysozyme (50 mg/mL) and 5 µL of RNase (1 mg/mL). The mixture was vortexed and incubated 30 min at 37°C. Twenty microliters of Protease K and 200 µL of Buffer AL were added. The mixture was vortexed again and incubated first during 30 min at 56°C, then for 5 min at 95°C. DNA was finally quantified in each individual extract using Quant-iT™ PicoGreen® dsDNA kit (Invitrogen; Thermo Fisher Scientific, Waltham, MA USA) and “Mock DNA” aliquots were prepared by mixing 2 µL of each extract, then stored at -80°C.

For the “Mock cloud” samples, the cell concentrations in individual cultures were estimated by flow cytometry. The six strains were then mixed at known concentrations (**Supplementary Table 1**) and 1 mL aliquots in 10% glycerol were stored at -80°C for further use. DNA was extracted in triplicate for mixed cell suspensions using either NucleoMag® DNA/RNA Water (Macherey-Nagel, Hoerdt, France), *i.e.*, the kit denominated as “Air” in the previous section, or QIAamp DNA Mini kit (QIAGEN; Hilden, Germany) (**Supplementary Figure 3**). DNA extracts were stored at -80°C before further processing. The relative abundances of 16S rRNA gene sequences in the extracts were estimated from total genomic DNA quantifications and from the mean numbers of ribosomes in the corresponding genera, as reported in the ribosomal RNA Database (rrnDB, v 5.7): 4.3 copies for *Rhodococcus*, 6.6 for *Flavobacterium*, 8.7 for *Bacillus*, 5.7 for *Staphylococcus*, 4.8 for *Pseudomonas*, and 2.0 for *Sphingomonas*.

### Bioinformatics data processing and statistics

Amplicon sequence variants (ASVs) were obtained from raw reads with the package *dada2* (v 1.20.0) (Callahan et al., 2016), using the functions *filterAndTrim*, *learnErrors*, *dada*, *mergePairs*, *makeSequenceTable* and *removeBimeraDenovo* following authors guidelines. Then, FROGS software (Bernard et al., 2021) was used to affiliate ASVs against SILVA v138.1 (Quast et al., 2013). When the BLAST assignation was questionable (*i.e.*, multi-affiliations, percent identity < 95 %, or percent query coverage < 98 %), this was verified using the RDP assignation and the EzBioCloud 16S rRNA gene-based ID database (Yoon et al., 2017; https://www.ezbiocloud.net/, update 2021.07.07). ASVs affiliated to *Chloroplast* (46 ASVs; 13% of total sequences), *Mitochondria* (71; 1.6 %) or Archaea (5; 0.2%), and ASVs without affiliation (87; 4%) were removed. The number of raw reads processed and remaining for analysis after curation and rarefaction are indicated in **Supplementary Table 2**. Environmental and mock samples were rarefied to 16,250 and 28,100 sequences, respectively, corresponding to the sample of each category with the lowest number of reads, using *FROGS Abundance normalization*. Blank samples were not rarefied. This left 22 and 556 ASVs in mock and environmental samples for further analyses, respectively.

ASV abundance data were centered log-ratio (CLR)-transformed, as recommended by Gloor et al. (2017) to account for their compositional nature. Data analysis was performed and represented using the *R* environment (v 4.0.3) (R Core Team (2019). R: A language and environment for statistical computing., 2020). The *zCompositions* package (v 1.3.4) (Palarea-Albaladejo and Martín-Fernández, 2015) was used to replace null counts in our compositional data based on a Bayesian-multiplicative method (function *cmultRepl* using CZM method and an input format in pseudo-counts) and to CLR-transform the abundance table (*clr* function). Heatmaps and dendograms were obtained using the packages *pheatmap* (v 1.0.12) (Raivo Kolde, 2019) and *ggdendro* (v 0.1.22) (Andrie de Vries and Brian D. Ripley, 2016). Principal component analysis was carried out using the R package *factoExtra,* with ellipses depicting 95% confidence levels. The R package *decontam* (Davis et al., 2018) was used for identifying and removing contaminants (referred to as “Method (*iv*)” in the corresponding result section). Univariate statistical tests were performed using PAST v. 4.02 (Hammer et al., 2001).

## Results and discussion

Total cells in the air samples averaged (1.98 ± 0.55)×10^3^ cell m^-3^ of air (**Table 1**); these are typical values at puy de Dôme station (Vaïtilingom et al., 2012). In total, 424 ASVs could be detected on Day 1, 378 on Day 2, and 348 on Day 3. The main bacterial phyla identified belonged to Proteobacteria (Alpha- and Gamma-; *e.g.*, *Sphingomonas*, *Methylobacterium, Massilia*, *Pseudomonas* and *Bradyrhizobium*), Firmicutes (*e.g.*, *Bacillus*), Actinobacteria (*e.g.*, *Nocardioides*), and Bacteroidota (*e.g.*, *Hymenobacter* and *Pedobacter*), which are commonly reported in the atmosphere (Bowers et al., 2012; Tignat-perrier et al., 2019). Day 1 was characterized by high abundance of *Sphingomonas* (3.2 % of the sequences), over *Hymenobacter* (1.1 %) and *Pseudomonas* (1.0 %), while Day 3 was more evenly dominated by *Sphingomonas* (1.8%), *Bacillus* (1.6 %) and *Methylobacterium* (1.2 %) (see **Supplementary Figure 4, and Supplementary Table 3**).

### The recovery of high amounts of DNA and RNA opens new perspectives

The amounts of DNA extracted from the collection liquids ranged from 70.5 ng (Day 3, replicate 2), to 324.5 ng (Day 1, replicate 3), with low variations between replicates (CV<30%). In total, 309 to 839 ng of DNA and 118.8 ng to 179.1 ng of RNA were available for analyses (**Figure 2**). These quantities are sufficient to allow direct sequencing of DNA and cDNA using current high throughput methods, which opens unprecedented opportunities of metagenomics and metatranscriptomics studies.

**Figure 2:**
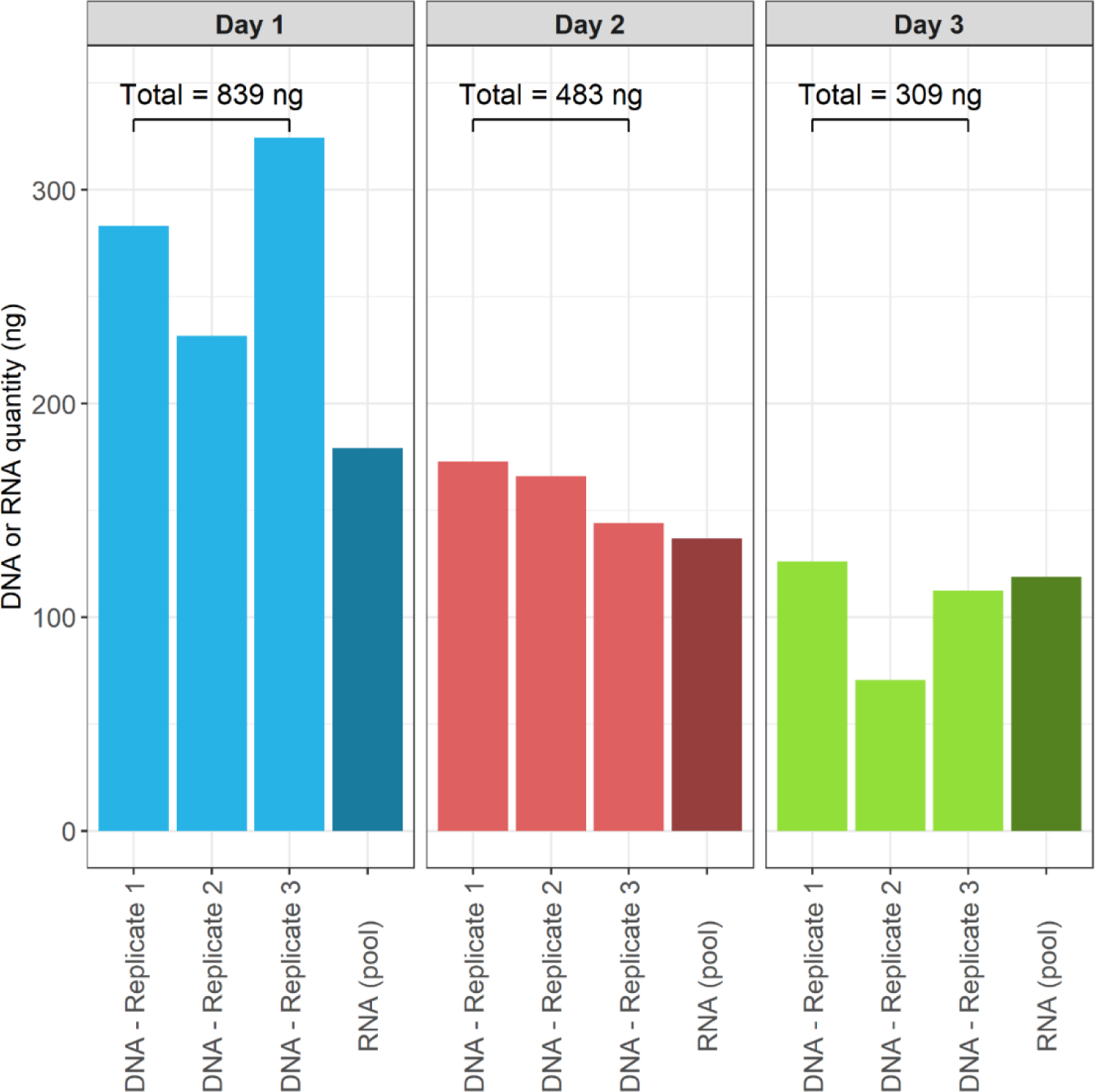
DNA and RNA concentrations in environmental air samples.

Metagenomics and metatranscriptomics approaches have proven their strength in the understanding of microbial functioning in environments like oceans (Salazar et al., 2019; Gifford et al., 2011). In the air, nucleic acids are in general obtained from filter samples (Be et al., 2015; Tignat-Perrier et al., 2020a; Gusareva et al., 2019), which prevents relevant assessment of transcriptomes. To our knowledge the very few studies reporting metatranscriptomes sequences from atmospheric samples so far involved targeted or untargeted pre-amplification (PCR or MDA) (Amato et al., 2019; Womack et al., 2015; Klein et al., 2016; Amato et al., 2017).

### Internal references allow evaluating quantitative aspects

Mixed cell suspensions (“Mock cloud”) and DNA mixes (“Mock DNA”), composed of cultures and DNA extracts of isolates from clouds, were processed as internal references. These included a range of bacteria typically reported in atmospheric samples: affiliated with *Pseudomonas syringae*, *Bacillus* sp., *Sphingomonas aerolata*, *Rhodococcus enclensis*, *Staphylococcus equorum* and *Flavobacterium* sp.. In all cases, clustering analysis followed expectations and grouped replicates with their respective theoretical taxa distribution (**Supplementary Figure 5**). Mock DNA samples data supported that PCR and sequencing processes were efficient at maintaining the relative distribution of taxa from DNA mixes. In Mock cloud samples, *Staphylococcus* was found overrepresented at the expense of *Bacillus* as compared to expectations based on cell counts, in particular when using QIAamp extraction kit. The primer pair used for 16S rRNA gene amplification (515F-806R) has been shown to perform better than many other current ribosomal primers in detecting a wide range of bacteria taxa (Abellan-Schneyder et al., 2021) and could not be incriminated. It is likely, rather, that cell counts by flow cytometry tended to underestimate the actual relative abundance of *Staphylococcus* in the original cell suspensions as this forms cell agglomerates, while *Bacillus* was to some extent recalcitrant to DNA extraction due to the presence of spores (Knüpfer et al., 2020). This indicates that specific taxa, in particular among Firmicutes, can present distortions in their relative abundance in sequence datasets compared with actual abundance in samples.

The few other sequences sporadically present in positive controls at very low abundance (<50 reads; <0.05% of the total reads in samples; **Supplementary Table 4**) may inform about the presence of contaminants and/or sequencing artifacts. Moreover, aliquots of both mock sample types were archived at -80°C for further use as references to compare data from distinct sequencing runs and studies.

### Replicate sampling allows statistics

The data demonstrate good reproducibility of the methods (**Figure 3**); it allowed distinguishing the airborne bacteria diversity of 3 consecutive days with very similar environmental conditions by principal component analysis (PCA) (**Figure 3A**).

**Figure 3:**
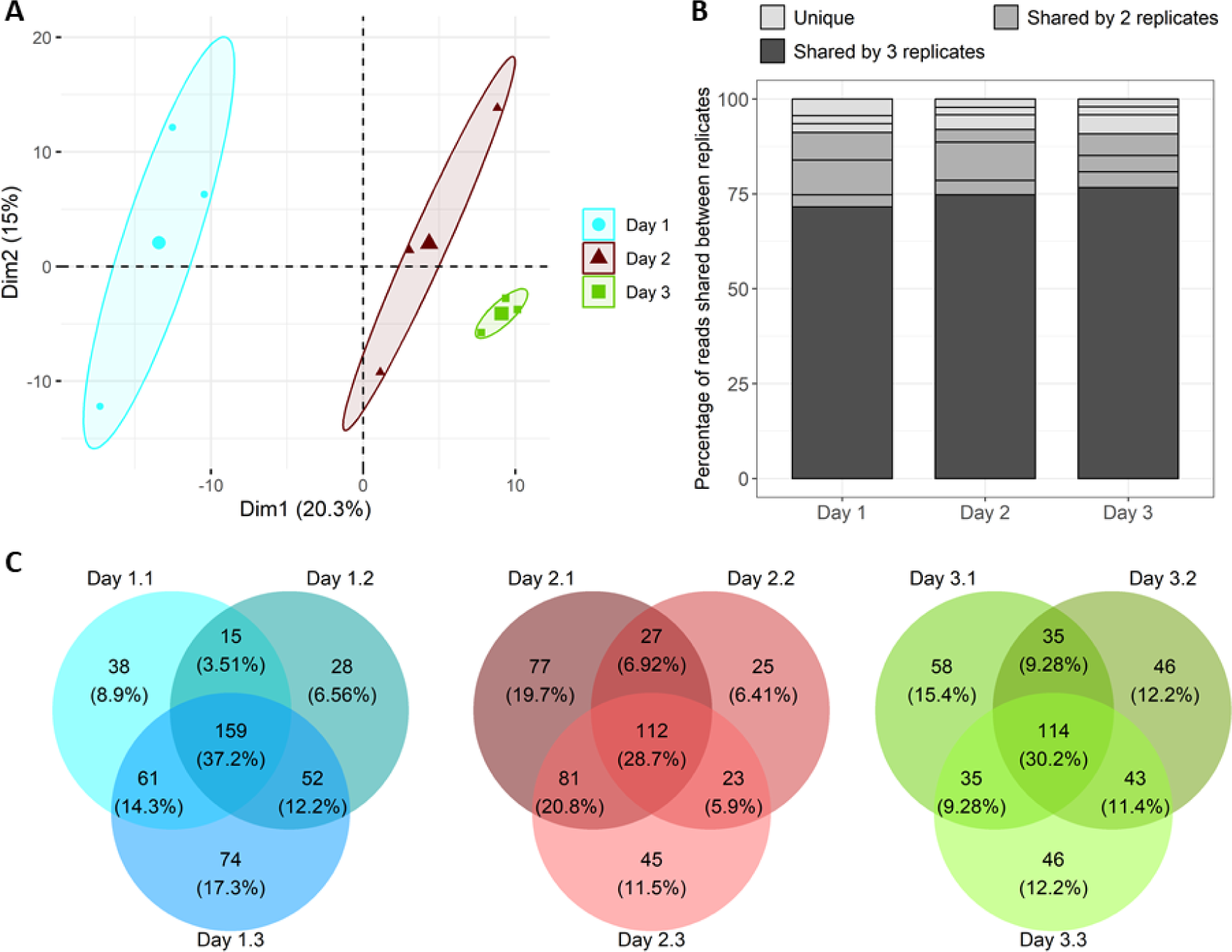
Distribution of ASVs and sequences among sample replicates. **A**: Principal component analysis based on ASV’s relative abundances (ellipses depict 95% confidence level areas); **B**: Proportions of sequencing reads retrieved in 1 (unique), 2 or 3 sample replicates; **C**: Venn diagrams showing the occurrence (presence/absence) of ASVs among sample replicates.

Bacterial assemblages were very uneven, composed of few abundant taxa accompanied by numerous sporadic (low-abundance) ones. Most sequences were consistently retrieved between sample replicates (71.6% to 76.8% depending on the sampling date; **Figure 3B**), representing a core assemblage gathering 28.7% to 37.2% of the richness (**Figure 3C**). In turn, only ∼3% of the sequences were specific, *i.e.* retrieved in only one of the 3 replicates of each sampling date, but they contributed relatively large proportions (6.4% to 19.7%) of the observed species richness in a sampler. Low-abundance taxa thus largely contributed to the variability observed between replicates and sampling dates, which is common in such datasets as pointed earlier in a range of ecosystems from air and lakes to skin and gut (Shade et al., 2014). With our setup, taxa’s rarity directly related to their chance to be captured by one or more samplers. The fact that a relatively large fraction of the total richness observed on a given sampling date was specific (∼33 to 40%) emphasizes the high spatial heterogeneity at short scale of the atmospheric environment, as regularly documented (Fierer et al., 2008), which is caused by a high proportion of rare taxa. The multiplicity of potential sources to the material collected and their respective strengths in emitting microorganisms can generate such variations despite overall very similar environmental conditions. Rare taxa can be important components of the material circulating in the air, for ecological and/or sanitary reasons, and they must not be neglected (Jalasvuori, 2020; Barberán et al., 2014; Leyronas et al., 2018; Pascoal et al., 2021; Pester et al., 2010; Rossi et al., 2022; Lynch and Neufeld, 2015).

Thanks to the individual processing of replicate samples, we could statistically discriminate sampling dates. Nevertheless, given the large number of potential explanatory variables involved respect to the limited number of samples assessed, it would be highly hazardous to try explaining the variability observed between sampling dates, so such attempt is not proposed. A more complete assessment of the extended dataset in relation to environmental variables can be found in Péguilhan et al. (2023).

### Account for the inevitable presence of contaminants

Processing negative controls is crucial in environmental microbiology to ensure data robustness, in particular where the biomass is low and subjected to strong short-scale variations such as in the atmosphere. As many contaminants can be brought by air itself (other than from the target environment) during sample handling and processing, detecting them is particularly challenging in aeromicrobiology (Šantl-Temkiv et al., 2022). Moreover, rare taxa can easily be confounded with stochastic trace contaminants from kits and reagents, or sequencing artefacts (de Goffau et al., 2018; Šantl-Temkiv et al., 2020; Glassing et al., 2016; Salter et al., 2014).

The amounts of nucleic acids recovered from controls, *i.e.* 1.7 L of unexposed deionized water (“water blanks”) or of NAP buffer exposed to the samplers (“sampling blanks”) were well below the amounts obtained from environmental samples: over 10 sampling blanks performed between July 2019 and September 2020 (including some which are not presented here, collected in the frame of other studies), the total amount of DNA ranged from 2.8 to 9.6 ng (7.45 ± 2.1 ng; mean ± SE), while it remained under detection limits for water blanks. From both control types, slight PCR products could be generated, and these were processed down to sequencing. This points toward the difficulty to maintain background signals at low level when using DNA amplification and high-throughput sequencing methods. Stinson et al. (2019) proposed to treat commercial reagents with DNase in order to remove potential contaminants.

The number of raw sequencing reads was much fewer in controls than in environmental samples as expected from lower amounts of DNA, with medians of ∼4,000 *versus* ∼88,000 reads, respectively (Mann-Whitney test, p = 0.001; **Supplementary Table 2**). Both water and sampling controls types exhibited much lower richness and diversity than environmental samples. Among controls, more raw reads could be obtained from unexposed water than from the collection liquid after exposed to the samplers, and they tended to be richer (Chao1 index). This confirms the effectiveness of sampler decontamination procedures and suggests that contamination occurred randomly during sample processing rather than during field work; it also indicates the absence of core contamination from the reagents and material used.

A total of 89 ASVs could be detected in controls, including 18 to 47 ASVs in water blanks and 24 to 29 ASVs in sampling blanks; 54 of them (∼61%) were also detected in environmental samples. These were affiliated with a large diversity of taxa identified as frequent contaminants in commercial reagents (Stinson et al., 2019; Salter et al., 2014; Glassing et al., 2016) and also typically dominant in atmospheric samples (Vaïtilingom et al., 2012; Amato et al., 2017), such *Sphingomonas*, *Pseudomonas*, *Hymenobacter*, and *Methylobacterium* (see **Supplementary Table 5** for complete list).

The stochasticity of contaminants (Stinson et al., 2019) makes them difficult to detect in complex datasets. In our study, most ASVs in controls (56 out of 89; 63%) occurred only once, and only 3 were recurring, affiliated with *Sphingomonas*, *Pseudomonas*, and *Burkholderia*: (cluster_4, cluster_2 and cluster_174, respectively). Cluster_4 was present in all the samples processed in this study, including positive (“Mock”) controls; cluster_2 was present in at least one replicate of each sampling date, in various proportions; cluster_174 was absent from environmental samples. This latter was therefore most likely associated with reagents or sequencing procedures.

The most conservative way to account for contaminants in samples is to exclude them totally from environmental datasets, regardless of their relative abundance (it would be irrelevant to subtract read numbers or proportions), although this is likely to remove also true members of the environment studied. Besides, less drastic, more elaborated statistical procedures have been proposed to account for both the prevalence and frequency of contaminants in such datasets (Davis et al., 2018), and so better consider stochasticity aspects; this requires replicates. In order to test the influence of sequence decontamination methods on richness and diversity, 4 treatments were applied to non-rarefied environmental datasets: (*i*) strict removal of all the ASVs detected in water blanks regardless of their relative abundance; (*ii*) strict removal of all the ASVs detected in sampling blanks; (*iii*) strict removal of all the ASVs detected in water and/or in sampling blanks; (*iv*) probability-based method cumulating the frequency and prevalence methods of the R package *decontam* (Davis et al., 2018).

The proportions of reads and ASVs removed with each method and their influence on richness and biodiversity on environmental data are presented in **Table 2** and **Figure 4**, respectively. All methods showed very good consistency between replicates. These all necessarily decreased richness in samples, by ∼5% to ∼13% in average depending on the methods (**Table 2**). The strict methods (*i*, *ii* and *iii*) led to the removal of higher proportions of reads than ASV, contrarily to the statistic method, and strongly increased biodiversity index in samples, by up to more than 1 point in the cases of Day 2 and Day 3 (**Figure 4**). In turn, the statistic method (*iv*) did not alter biodiversity indexes.

**Figure 4:**
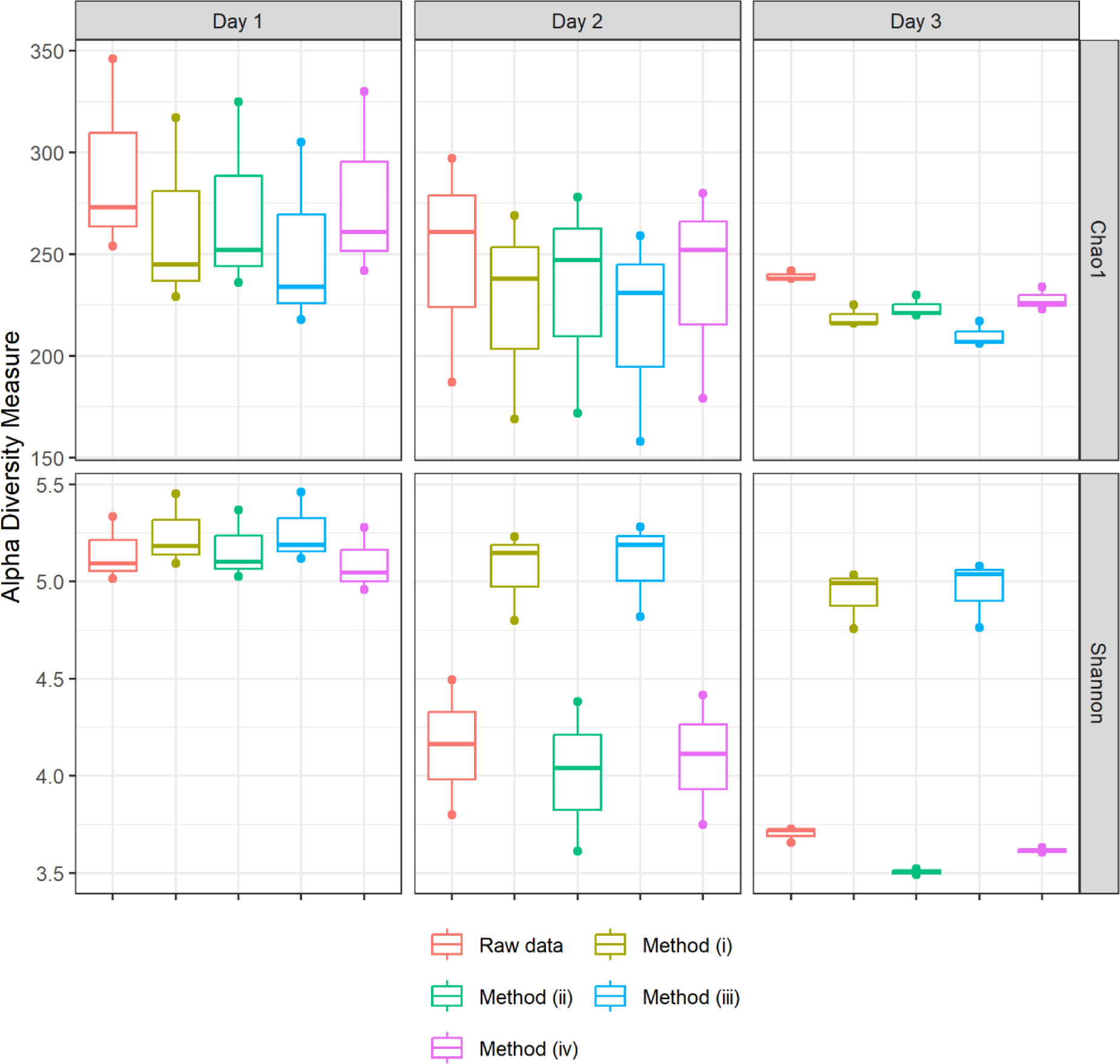
Influence of decontamination methods on environmental samples’ richness and biodiversity indexes.

**Table 2:** Proportions of reads and ASVs removed from environmental datasets from decontamination methods (see text for details on the methods).

|  | **Initial data (non-rarefied)** |  | **Method (i) (water blanks)** |  | **Method (ii) (sampling blanks)** |  | **Method (iii) (water + sampling blanks)** |  | **Method (iv) (probability)** |  |
| --- | --- | --- | --- | --- | --- | --- | --- | --- | --- | --- |
| **Sample** | Number of reads | Number of ASVs | % reads removed | % ASVs removed | % reads removed | % ASVs removed | % reads removed | % ASVs removed | % reads removed | % ASVs removed |
| Day 1.1 | 18823 | 273 | 24.07% | 10.26% | 16.66% | 7.69% | 31.07% | 14.29% | 2.75% | 4.40% |
| Day 1.2 | 16250 | 254 | 23.22% | 9.84% | 16.46% | 7.09% | 31.41% | 14.17% | 3.24% | 4.72% |
| Day 1.3 | 24856 | 346 | 22.55% | 8.38% | 14.97% | 6.07% | 29.03% | 11.85% | 2.99% | 4.62% |
| Day 2.1 | 27670 | 297 | 38.23% | 9.43% | 12.94% | 6.40% | 44.86% | 12.79% | 2.97% | 5.72% |
| Day 2.2 | 19027 | 187 | 45.99% | 9.63% | 12.66% | 8.02% | 53.82% | 15.51% | 1.56% | 4.28% |
| Day 2.3 | 24406 | 261 | 44.14% | 8.81% | 12.27% | 5.36% | 49.63% | 11.49% | 1.63% | 3.45% |
| Day 3.1 | 29709 | 242 | 45.88% | 7.02% | 9.41% | 4.96% | 51.68% | 10.33% | 1.50% | 3.31% |
| Day 3.2 | 24392 | 238 | 48.38% | 9.24% | 11.30% | 7.56% | 55.12% | 13.45% | 3.23% | 6.30% |
| Day 3.3 | 23014 | 238 | 48.94% | 9.24% | 10.72% | 7.14% | 54.94% | 13.03% | 2.92% | 5.04% |
| **Mean** |  |  | **37.93%** | **9.10%** | **13.04%** | **6.70%** | **44.62%** | **12.99%** | **2.53%** | **4.65%** |
| *Standard error* |  |  | *11.41%* | *0.95%* | *2.52%* | *1.07%* | *11.05%* | *1.59%* | *0.74%* | *0.97%* |

The decontamination solutions that can appear the most conservative at first sight can thus counterintuitively lead to increase biodiversity in environmental samples and smooth temporal variability, so result in erroneous conclusions. In addition of strengthening statistics aiming at deciphering alpha- and beta-diversity of airborne microbial assemblages, replication also enables more meaningful decontamination methods to be applied.

## Conclusion

We used an analytical framework involving some of the basic recommended practices in environmental microbiology (replicate, check and account for contaminants, include internal references) to atmospheric samples, and demonstrate that these can logically benefit to aerobiology as well in many aspects. Replicated sampling at high air-flow rate for short periods of time into nucleic acid preservation buffer allows capturing close atmospheric situations that can be distinguished statistically using high throughput sequencing methods. Replication is also necessary to account for the stochastic and inevitable presence of contaminants, and so to ensure confidence to taxa’s presence and abundance in environmental samples. Finally, replicating also allows pooling to access larger amounts of material if necessary, which opens new perspectives for metagenomics and metatranscriptomics investigations of the aeromicrobiome. Analyses of the alpha- and beta-diversity of microbial assemblages in air masse, and short-scale and specific atmospheric variations (*i.e.*, day-to-day or day/night variations, clouds, pollution events, etc) will be assessable with good confidence using this framework.

## Author contributions

RP, MJ have made major contribution to the acquisition, analysis and interpretation of the data. FR and OR helped with data acquisition and data analysis. PA conceived and designed the study, and interpreted the data. PA and RP wrote the manuscript.

## Data Archiving Statement

Demultiplexed sequencing files were deposited at the European Nucleotide Archive and have accession numbers ERR9924984 to ERR9924999, and ERR9924950 to ERR9924958 (project accession: PRJEB54614).

## Acknowledgments

We thank OPGC’ staff for access to puy de Dôme instrumented station. We are grateful to the platforms AUBI (Clermont Auvergne University’s Mésocentre) for providing computing and storage capacities, and CYSTEM (UCA Partner service) for access to flow cytometry. This research was supported by the French National Research Agency (ANR) (grant no. ANR-17-MPGA-0013), with fellowship from Clermont Auvergne University to RP.

## References

Abellan-Schneyder, I., Matchado, M. S., Reitmeier, S., Sommer, A., Sewald, Z., Baumbach, J., List, M., and Neuhaus, K.: Primer, Pipelines, Parameters: Issues in 16S rRNA Gene Sequencing, mSphere, 6, e01202–20, 10.1128/mSphere.01202-20, 2021.

Amato, P., Parazols, M., Sancelme, M., Laj, P., Mailhot, G., and Delort, A. M.: Microorganisms isolated from the water phase of tropospheric clouds at the Puy de Dôme: Major groups and growth abilities at low temperatures, in: FEMS Microbiology Ecology, 242–254, 10.1111/j.1574-6941.2006.00199.x, 2007.

Amato, P., Joly, M., Besaury, L., Oudart, A., Taib, N., Moné, A. I., Deguillaume, L., Delort, A. M., and Debroas, D.: Active microorganisms thrive among extremely diverse communities in cloud water, PLoS ONE, 12, 1–22, 10.1371/journal.pone.0182869, 2017.

Amato, P., Besaury, L., Joly, M., Penaud, B., Deguillaume, L., and Delort, A. M.: Metatranscriptomic exploration of microbial functioning in clouds, Scientific Reports, 9, 10.1038/s41598-019-41032-4, 2019.

Andrie de Vries and Brian D. Ripley: ggdendro: Create Dendrograms and Tree Diagrams Using “ggplot2”. R package version 0.1-20., 2016.

Apprill, A., Mcnally, S., Parsons, R., and Weber, L.: Minor revision to V4 region SSU rRNA 806R gene primer greatly increases detection of SAR11 bacterioplankton, Aquatic Microbial Ecology, 75, 129–137, 10.3354/ame01753, 2015.

Archer, S. D. J., Lee, K. C., Caruso, T., Maki, T., Lee, C. K., Cary, S. C., Cowan, D. A., Maestre, F. T., and Pointing, S. B.: Airborne microbial transport limitation to isolated Antarctic soil habitats, Nature Microbiology, 4, 925–932, 10.1038/s41564-019-0370-4, 2019.

Baray, J. L., Deguillaume, L., Colomb, A., Sellegri, K., Freney, E., Rose, C., Baelen, J. Van, Pichon, J. M., Picard, D., Fréville, P., Bouvier, L., Ribeiro, M., Amato, P., Banson, S., Bianco, A., Borbon, A., Bourcier, L., Bras, Y., Brigante, M., Cacault, P., Chauvigne, A., Charbouillot, T., Chaumerliac, N., Delort, A. M., Delmotte, M., Dupuy, R., Farah, A., Febvre, G., Flossmann, A., Gourbeyre, C., Hervier, C., Hervo, M., Huret, N., Joly, M., Kazan, V., Lopez, M., Mailhot, G., Marinoni, A., Masson, O., Montoux, N., Parazols, M., Peyrin, F., Pointin, Y., Ramonet, M., Rocco, M., Sancelme, M., Sauvage, S., Schmidt, M., Tison, E., Vaïtilingom, M., Villani, P., Wang, M., Yver-Kwok, C., and Laj, P.: Cézeaux-Aulnat-Opme-Puy de Dôme: A multi-site for the long-term survey of the tropospheric composition and climate change, Atmospheric Measurement Techniques, 13, 3413–3445, 10.5194/amt-13-3413-2020, 2020.

Barberán, A., Henley, J., Fierer, N., and Casamayor, E. O.: Structure, inter-annual recurrence, and global-scale connectivity of airborne microbial communities, Sci. Total Environ., 487, 187–195, 10.1016/j.scitotenv.2014.04.030, 2014.

Barberán, A., Ladau, J., Leff, J. W., Pollard, K. S., Menninger, H. L., Dunn, R. R., and Fierer, N.: Continental-scale distributions of dust-associated bacteria and fungi, PNAS, 112, 5756– 5761, 10.1073/pnas.1420815112, 2015.

Bauer, H., Giebl, H., Hitzenberger, R., Kasper-Giebl, A., Reischl, G., Zibuschka, F., and Puxbaum, H.: Airborne bacteria as cloud condensation nuclei, Journal of Geophysical Research: Atmospheres, 108, 4658, 10.1029/2003JD003545, 2003.

Be, N. A., Thissen, J. B., Fofanov, V. Y., Allen, J. E., Rojas, M., Golovko, G., Fofanov, Y., Koshinsky, H., and Jaing, C. J.: Metagenomic analysis of the airborne environment in urban spaces, Microb. Ecol., 69, 346–355, 10.1007/s00248-014-0517-z, 2015.

Beattie, G. A. and Lindow, S. E.: The secret life of foliar bacterial pathogens on leaves, Annual review of phytopathology, 33, 145–172, 1995.

Bernard, M., Rué, O., Mariadassou, M., and Pascal, G.: FROGS: a powerful tool to analyse the diversity of fungi with special management of internal transcribed spacers, Briefings in Bioinformatics, 22, 10.1093/BIB/BBAB318, 2021.

Bowers, R. M., McCubbin, I. B., Hallar, A. G., and Fierer, N.: Seasonal variability in airborne bacterial communities at a high-elevation site, Atmospheric Environment, 50, 41–49, 10.1016/j.atmosenv.2012.01.005, 2012.

Burrows, S. M., Elbert, W., Lawrence, M. G., and Pöschl, U.: Bacteria in the global atmosphere – Part 1: Review and synthesis of literature data for different ecosystems, Atmospheric Chemistry and Physics, 9, 9263–9280, 10.5194/acp-9-9263-2009, 2009.

Callahan, B. J., McMurdie, P. J., Rosen, M. J., Han, A. W., Johnson, A. J. A., and Holmes, S. P.: DADA2: High-resolution sample inference from Illumina amplicon data, Nature Methods 2016 13:7, 13, 581–583, 10.1038/nmeth.3869, 2016.

Camacho-Sanchez, M., Burraco, P., Gomez-Mestre, I., and Leonard, J. A.: Preservation of RNA and DNA from mammal samples under field conditions, Molecular Ecology Resources, 13, 663–673, 10.1111/1755-0998.12108, 2013.

Davis, N. M., Proctor, D. M., Holmes, S. P., Relman, D. A., and Callahan, B. J.: Simple statistical identification and removal of contaminant sequences in marker-gene and metagenomics data, Microbiome, 6, 226, 10.1186/s40168-018-0605-2, 2018.

Després, V. R., Alex Huffman, J., Burrows, S. M., Hoose, C., Safatov, A. S., Buryak, G., Fröhlich-Nowoisky, J., Elbert, W., Andreae, M. O., Pöschl, U., and Jaenicke, R.: Primary biological aerosol particles in the atmosphere: A review, Tellus, Series B: Chemical and Physical Meteorology, 64, 10.3402/tellusb.v64i0.15598, 2012.

Dommergue, A., Amato, P., Tignat-Perrier, R., Magand, O., Thollot, A., Joly, M., Bouvier, L., Sellegri, K., Vogel, T., Sonke, J. E., Jaffrezo, J. L., Andrade, M., Moreno, I., Labuschagne, C., Martin, L., Zhang, Q., and Larose, C.: Methods to investigate the global atmospheric microbiome, Frontiers in Microbiology, 10, 10.3389/fmicb.2019.00243, 2019.

Dybwad, M., Skogan, G., and Blatny, J. M.: Comparative Testing and Evaluation of Nine Different Air Samplers: End-to-End Sampling Efficiencies as Specific Performance Measurements for Bioaerosol Applications, Aerosol Science and Technology, 48, 282–295, 10.1080/02786826.2013.871501, 2014.

Fernstrom, A., Goldblatt, M., Fernstrom, A., and Goldblatt, M.: Aerobiology and Its Role in the Transmission of Infectious Diseases, Aerobiology and Its Role in the Transmission of Infectious Diseases, Journal of Pathogens, Journal of Pathogens, 2013, 2013, e493960, 10.1155/2013/493960, 2013.

Fierer, N., Liu, Z., Rodríguez-Hernández, M., Knight, R., Henn, M., and Hernandez, M. T.: Short-Term Temporal Variability in Airborne Bacterial and Fungal Populations, Appl Environ Microbiol, 74, 200–207, 10.1128/AEM.01467-07, 2008.

Gifford, S. M., Sharma, S., Rinta-Kanto, J. M., and Moran, M. A.: Quantitative analysis of a deeply sequenced marine microbial metatranscriptome, The ISME Journal, 5, 461–472, 10.1038/ismej.2010.141, 2011.

Glassing, A., Dowd, S. E., Galandiuk, S., Davis, B., and Chiodini, R. J.: Inherent bacterial DNA contamination of extraction and sequencing reagents may affect interpretation of microbiota in low bacterial biomass samples, Gut Pathog, 8, 24, 10.1186/s13099-016-0103-7, 2016.

de Goffau, M. C., Lager, S., Salter, S. J., Wagner, J., Kronbichler, A., Charnock-Jones, D. S., Peacock, S. J., Smith, G. C. S., and Parkhill, J.: Recognizing the reagent microbiome, Nature Microbiology, 3, 851–853, 10.1038/s41564-018-0202-y, 2018.

Griffin, D. W., Gonzalez, C., Teigell, N., Petrosky, T., Northup, D. E., and Lyles, M.: Observations on the use of membrane filtration and liquid impingement to collect airborne microorganisms in various atmospheric environments, Aerobiologia, 27, 25–35, 10.1007/s10453-010-9173-z, 2011.

Gusareva, E. S., Acerbi, E., Lau, K. J. X., Luhung, I., Premkrishnan, B. N. V., Kolundžija, S., Purbojati, R. W., Wong, A., Houghton, J. N. I., Miller, D., Gaultier, N. E., Heinle, C. E., Clare, M. E., Vettath, V. K., Kee, C., Lim, S. B. Y., Chénard, C., Phung, W. J., Kushwaha, K. K., Nee, A. P., Putra, A., Panicker, D., Yanqing, K., Hwee, Y. Z., Lohar, S. R., Kuwata, M., Kim, H. L., Yang, L., Uchida, A., Drautz-Moses, D. I., Junqueira, A. C. M., and Schuster, S. C.: Microbial communities in the tropical air ecosystem follow a precise diel cycle, Proceedings of the National Academy of Sciences, 116, 23299–23308, 10.1073/pnas.1908493116, 2019.

Hammer, Ø., Harper, D. A. T., and Ryan, P. D.: Past: Paleontological statistics software package for education and data analysis, Palaeontologia Electronica, 4, 1–9, 2001.

Jalasvuori, M.: Silent rain: does the atmosphere-mediated connectivity between microbiomes influence bacterial evolutionary rates?, FEMS Microbiol Ecol, 96, 10.1093/femsec/fiaa096, 2020.

Ji, B. W., Sheth, R. U., Dixit, P. D., Huang, Y., Kaufman, A., Wang, H. H., and Vitkup, D.: Quantifying spatiotemporal variability and noise in absolute microbiota abundances using replicate sampling, Nat Methods, 16, 731–736, 10.1038/s41592-019-0467-y, 2019.

Joly, M., Amato, P., Sancelme, M., Vinatier, V., Abrantes, M., Deguillaume, L., and Delort, A.-M.: Survival of microbial isolates from clouds toward simulated atmospheric stress factors, Atmospheric Environment, 117, 92–98, 10.1016/j.atmosenv.2015.07.009, 2015.

Kathiriya, T., Gupta, A., and Singh, N. K.: An opinion review on sampling strategies, enumeration techniques, and critical environmental factors for bioaerosols: An emerging sustainability indicator for society and cities, 10.1016/j.eti.2020.101287, 2021.

Khaled, A., Zhang, M., Amato, P., Delort, A. M., and Ervens, B.: Biodegradation by bacteria in clouds: An underestimated sink for some organics in the atmospheric multiphase system, Atmospheric Chemistry and Physics, 21, 3123–3141, 10.5194/acp-21-3123-2021, 2021.

Klein, A. M., Bohannan, B. J. M., Jaffe, D. A., Levin, D. A., and Green, J. L.: Molecular Evidence for Metabolically Active Bacteria in the Atmosphere, Front. Microbiol., 772, 10.3389/fmicb.2016.00772, 2016.

Knüpfer, M., Braun, P., Baumann, K., Rehn, A., Antwerpen, M., Grass, G., and Wölfel, and R.: Evaluation of a Highly Efficient DNA Extraction Method for Bacillus anthracis Endospores, Microorganisms, 8, 763, 10.3390/microorganisms8050763, 2020.

Krumins, V., Mainelis, G., Kerkhof, L. J., and Fennell, D. E.: Substrate-Dependent rRNA Production in an Airborne Bacterium, Environmental Science and Technology Letters, 1, 376–381, 10.1021/ez500245y, 2014.

Lallement, A., Besaury, L., Eyheraguibel, B., Amato, P., Sancelme, M., Mailhot, G., and Delort, A. M.: Draft Genome Sequence of Rhodococcus enclensis 23b-28, a Model Strain Isolated from Cloud Water, Genome Announcements, 5, 10.1128/genomea.01199-17, 2017.

Leyronas, C., Morris, C. E., Choufany, M., and Soubeyrand, S.: Assessing the Aerial Interconnectivity of Distant Reservoirs of Sclerotinia sclerotiorum, Front. Microbiol., 9, 10.3389/fmicb.2018.02257, 2018.

Lynch, M. D. J. and Neufeld, J. D.: Ecology and exploration of the rare biosphere, Nat Rev Microbiol, 13, 217–229, 10.1038/nrmicro3400, 2015.

Manibusan, S. and Mainelis, G.: Passive bioaerosol samplers: A complementary tool for bioaerosol research. A review, Journal of Aerosol Science, 163, 105992, 10.1016/j.jaerosci.2022.105992, 2022.

Möhler, O., DeMott, P. J., Vali, G., and Levin, Z.: Microbiology and atmospheric processes: The role of biological particles in cloud physics, 10.5194/bg-4-1059-2007, 2007.

Palarea-Albaladejo, J. and Martín-Fernández, J. A.: ZCompositions - R package for multivariate imputation of left-censored data under a compositional approach, Chemometrics and Intelligent Laboratory Systems, 143, 85–96, 10.1016/j.chemolab.2015.02.019, 2015.

Parada, A. E., Needham, D. M., and Fuhrman, J. A.: Every base matters: Assessing small subunit rRNA primers for marine microbiomes with mock communities, time series and global field samples, Environmental Microbiology, 18, 1403–1414, 10.1111/1462-2920.13023, 2016.

Pascoal, F., Costa, R., and Magalhães, C.: The microbial rare biosphere: current concepts, methods and ecological principles, FEMS Microbiol Ecol, 97, fiaa227, 10.1093/femsec/fiaa227, 2021.

Patade, S., Phillips, V. T. J., Amato, P., Bingemer, H. G., Burrows, S. M., DeMott, P. J., Goncalves, F. L. T., Knopf, D. A., Morris, C. E., Alwmark, C., Artaxo, P., Pöhlker, C., Schrod, J., and Weber, B.: Empirical formulation for multiple groups of primary biological ice nucleating particles from field observations over Amazonia, Journal of the Atmospheric Sciences, 1, 10.1175/JAS-D-20-0096.1, 2021.

Péguilhan, R., Besaury, L., Rossi, F., Enault, F., Baray, J., Deguillaume, L., and Amato, P.: Rainfalls sprinkle cloud bacterial diversity while scavenging biomass, FEMS Microbiology Ecology, 1–15, 10.1093/femsec/fiab144, 2021.

Péguilhan, R., Rossi, F., Rué, O., Joly, M., and Amato, P.: Comparative analysis of bacterial diversity in clouds and aerosols, Atmospheric Environment, 119635, 10.1016/j.atmosenv.2023.119635, 2023.

Pester, M., Bittner, N., Deevong, P., Wagner, M., and Loy, A.: A ‘rare biosphere’ microorganism contributes to sulfate reduction in a peatland, ISME J, 4, 1591–1602, 10.1038/ismej.2010.75, 2010.

Prosser, J. I.: Replicate or lie, Environmental Microbiology, 12, 1806–1810, 10.1111/j.1462-2920.2010.02201.x, 2010.

Quast, C., Pruesse, E., Yilmaz, P., Gerken, J., Schweer, T., Yarza, P., Peplies, J., and Glöckner, F. O.: The SILVA ribosomal RNA gene database project: Improved data processing and web-based tools, Nucleic Acids Research, 41, 590–596, 10.1093/nar/gks1219, 2013.

R Core Team (2019). R: A language and environment for statistical computing.: http://www.r-project.org/index.html, last access: 13 April 2020.

Raivo Kolde: pheatmap: Pretty Heatmaps. R package version 1.0.12., 2019.

Rossi, F., Péguilhan, R., Turgeon, N., Veillette, M., Baray, J.-L., Deguillaume, L., Amato, P., and Duchaine, C.: Quantification of antibiotic resistance genes (ARGs) in clouds at a mountain site (puy de Dôme, central France), Science of The Total Environment, 161264, 10.1016/j.scitotenv.2022.161264, 2022.

Rule, A. M., Kesavan, J., Schwab, K. J., and Buckley, T. J.: Application of Flow Cytometry for the Assessment of Preservation and Recovery Efficiency of Bioaerosol Samplers Spiked with Pantoea agglomerans, Environ. Sci. Technol., 41, 2467–2472, 10.1021/es062394l, 2007.

Salazar, G., Paoli, L., Alberti, A., Huerta-Cepas, J., Ruscheweyh, H.-J., Cuenca, M., Field, C. M., Coelho, L. P., Cruaud, C., Engelen, S., Gregory, A. C., Labadie, K., Marec, C., Pelletier, E., Royo-Llonch, M., Roux, S., Sánchez, P., Uehara, H., Zayed, A. A., Zeller, G., Carmichael, M., Dimier, C., Ferland, J., Kandels, S., Picheral, M., Pisarev, S., Poulain, J., Acinas, S. G., Babin, M., Bork, P., Boss, E., Bowler, C., Cochrane, G., de Vargas, C., Follows, M., Gorsky, G., Grimsley, N., Guidi, L., Hingamp, P., Iudicone, D., Jaillon, O., Kandels-Lewis, S., Karp-Boss, L., Karsenti, E., Not, F., Ogata, H., Pesant, S., Poulton, N., Raes, J., Sardet, C., Speich, S., Stemmann, L., Sullivan, M. B., Sunagawa, S., Wincker, P., Acinas, S. G., Babin, M., Bork, P., Bowler, C., de Vargas, C., Guidi, L., Hingamp, P., Iudicone, D., Karp-Boss, L., Karsenti, E., Ogata, H., Pesant, S., Speich, S., Sullivan, M. B., Wincker, P., and Sunagawa, S.: Gene Expression Changes and Community Turnover Differentially Shape the Global Ocean Metatranscriptome, Cell, 179, 1068–1083.e21, 10.1016/j.cell.2019.10.014, 2019.

Salter, S. J., Cox, M. J., Turek, E. M., Calus, S. T., Cookson, W. O., Moffatt, M. F., Turner, P., Parkhill, J., Loman, N. J., and Walker, A. W.: Reagent and laboratory contamination can critically impact sequence-based microbiome analyses, BMC Biology, 12, 87, 10.1186/s12915-014-0087-z, 2014.

Šantl-Temkiv, T., Amato, P., Gosewinkel, U., Thyrhaug, R., Charton, A., Chicot, B., Finster, K., Bratbak, G., and Löndahl, J.: High-Flow-Rate Impinger for the Study of Concentration, Viability, Metabolic Activity, and Ice-Nucleation Activity of Airborne Bacteria, Environmental Science and Technology, 51, 11224–11234, 10.1021/acs.est.7b01480, 2017.

Šantl-Temkiv, T., Sikoparija, B., Maki, T., Carotenuto, F., Amato, P., Yao, M., Morris, C. E., Schnell, R., Jaenicke, R., Pöhlker, C., DeMott, P. J., Hill, T. C. J., and Huffman, J. A.: Bioaerosol field measurements: Challenges and perspectives in outdoor studies, Aerosol Science and Technology, 54, 520–546, 10.1080/02786826.2019.1676395, 2020.

Šantl-Temkiv, T., Amato, P., Casamayor, E. O., Lee, P. K. H., and Pointing, S. B.: Microbial ecology of the atmosphere, FEMS Microbiology Reviews, fuac009, 10.1093/femsre/fuac009, 2022.

Shade, A., Jones, S. E., Caporaso, J. G., Handelsman, J., Knight, R., Fierer, N., and Gilbert, J. A.: Conditionally Rare Taxa Disproportionately Contribute to Temporal Changes in Microbial Diversity, mBio, 5, e01371–14, 10.1128/mBio.01371-14, 2014.

Smith, D. J., Timonen, H. J., Jaffe, D. A., Griffin, D. W., Birmele, M. N., Perry, K. D., Ward, P. D., and Roberts, M. S.: Intercontinental dispersal of bacteria and archaea by transpacific winds, Applied and Environmental Microbiology, 79, 1134–1139, 10.1128/AEM.03029-12, 2013.

Stinson, L. F., Keelan, J. A., and Payne, M. S.: Identification and removal of contaminating microbial DNA from PCR reagents: impact on low-biomass microbiome analyses, Letters in Applied Microbiology, 68, 2–8, 10.1111/lam.13091, 2019.

Tignat-perrier, R., Dommergue, A., Thollot, A., Keuschnig, C., Magand, O., Vogel, T. M., and Larose, C.: Global airborne microbial communities controlled by surrounding landscapes and wind conditions, Scientific Reports, 1–11, 10.1038/s41598-019-51073-4, 2019.

Tignat-Perrier, R., Dommergue, A., Thollot, A., Magand, O., Vogel, T. M., and Larose, C.: Microbial functional signature in the atmospheric boundary layer, Biogeosciences, 17, 6081– 6095, 10.5194/bg-17-6081-2020, 2020a.

Tignat-Perrier, R., Dommergue, A., Thollot, A., Magand, O., Amato, P., Joly, M., Sellegri, K., Vogel, T. M., and Larose, C.: Seasonal shift in airborne microbial communities, Science of The Total Environment, 137129, 10.1016/j.scitotenv.2020.137129, 2020b.

Vaïtilingom, M., Attard, E., Gaiani, N., Sancelme, M., Deguillaume, L., Flossmann, A. I., Amato, P., and Delort, A. M.: Long-term features of cloud microbiology at the puy de Dôme (France), Atmospheric Environment, 56, 88–100, 10.1016/j.atmosenv.2012.03.072, 2012.

Wirgot, N., Vinatier, V., Deguillaume, L., Sancelme, M., and Delort, A. M.: H2O2 modulates the energetic metabolism of the cloud microbiome, Atmospheric Chemistry and Physics, 17, 14841–14851, 10.5194/acp-17-14841-2017, 2017.

Womack, A. M., Artaxo, P. E., Ishida, F. Y., Mueller, R. C., Saleska, S. R., Wiedemann, K. T., Bohannan, B. J. M., and Green, J. L.: Characterization of active and total fungal communities in the atmosphere over the Amazon rainforest, Biogeosciences, 12, 6337–6349, 10.5194/bg-12-6337-2015, 2015.

Yoon, S. H., Ha, S. M., Kwon, S., Lim, J., Kim, Y., Seo, H., and Chun, J.: Introducing EzBioCloud: A taxonomically united database of 16S rRNA gene sequences and whole-genome assemblies, International Journal of Systematic and Evolutionary Microbiology, 67, 1613–1617, 10.1099/ijsem.0.001755, 2017.

Zhang, M., Khaled, A., Amato, P., Delort, A. M., and Ervens, B.: Sensitivities to biological aerosol particle properties and ageing processes: Potential implications for aerosol-cloud interactions and optical properties, Atmospheric Chemistry and Physics, 21, 3699–3724, 10.5194/acp-21-3699-2021, 2021.

Zhao, J., Jin, L., Wu, D., Xie, J., Li, J., Fu, X., Cong, Z., Fu, P., Zhang, Y., Luo, X., Feng, X., Zhang, G., Tiedje, J. M., and Li, X.: Global airborne bacterial community—interactions with Earth’s microbiomes and anthropogenic activities, Proceedings of the National Academy of Sciences, 119, e2204465119, 10.1073/pnas.2204465119, 2022.

