## Supplementary Figure 1 for "Experimental and methodological framework for the assessment of nucleic acids in airborne microorganisms"

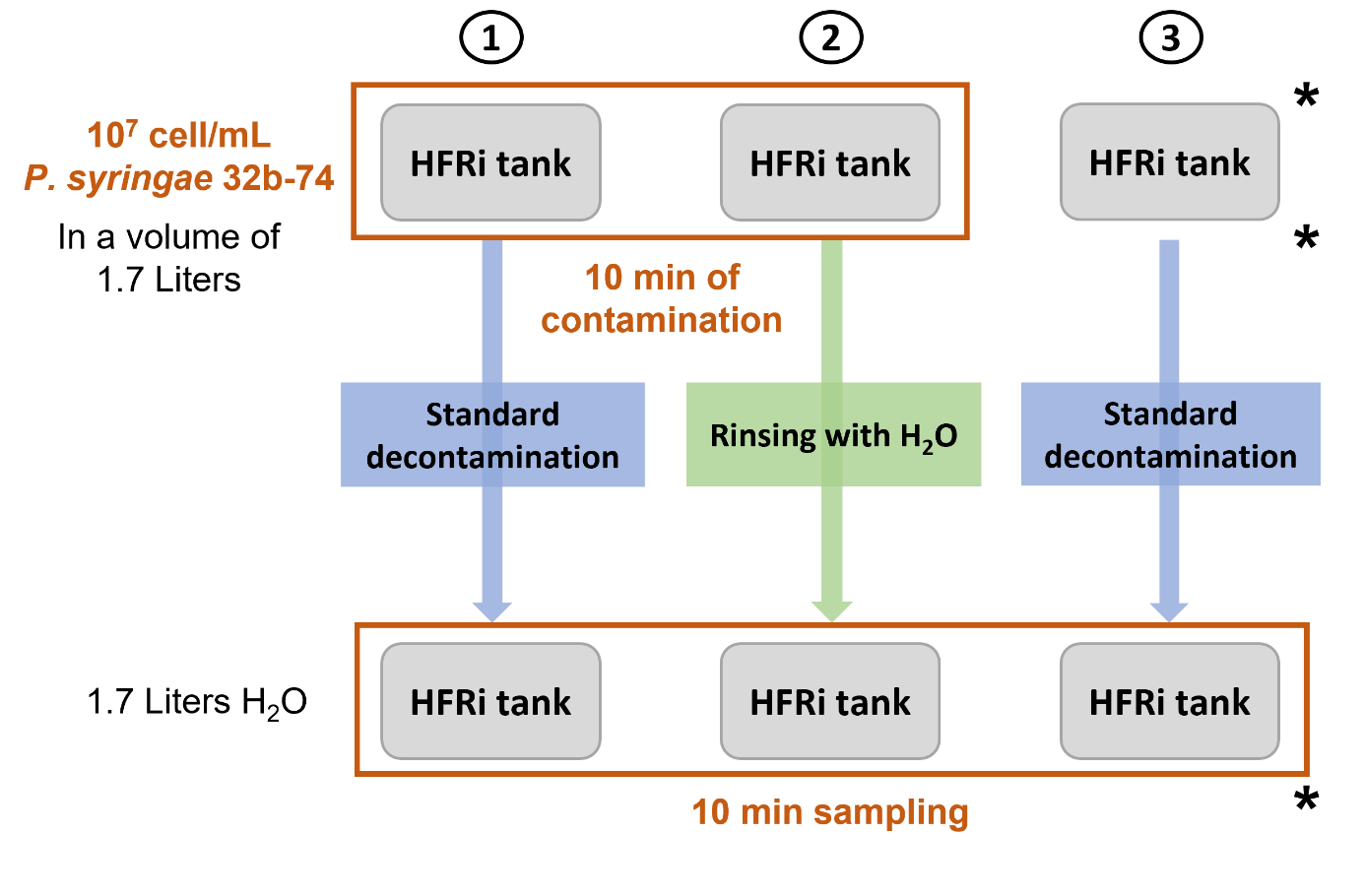

**Supplementary Figure 1:** Schematic of the protocol used for assessing sampler decontamination procedures. Asterisks indicate sampling steps for controlling the presence of contaminants.
