## Supplementary Figure 2 for "Experimental and methodological framework for the assessment of nucleic acids in airborne microorganisms"

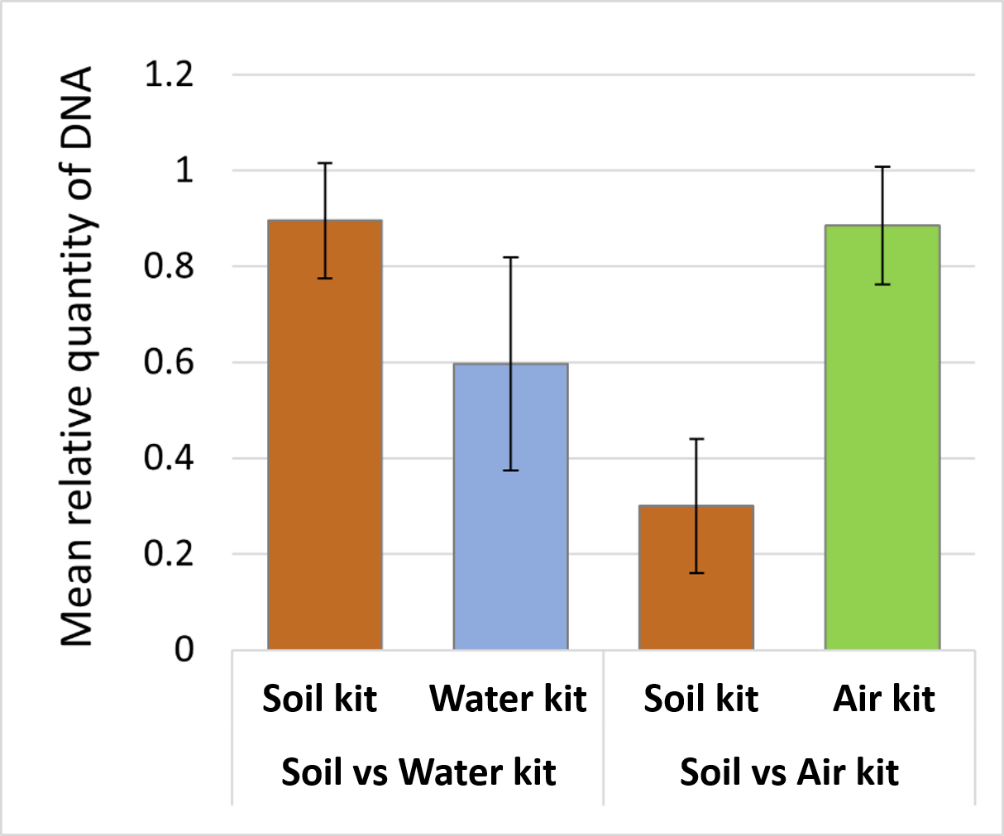

**Supplementary Figure 2:** Relative amount of DNA retrieved from air samples with the different extraction kits. Paired-comparison of three commercial nucleic acid extraction kits. “Water”: DNeasy PowerWater kit (QIAGEN; Hilden, Germany); “Soil”: NucleoSpin Soil kit (Macherey-Nagel, Hoerdt, France); “Air”: NucleoMag DNA/RNA Water kit (Macherey-Nagel, Hoerdt, France). N = 2 sampling dates for each paired test.
