## Supplementary figures and images for "Experimental and methodological framework for the assessment of nucleic acids in airborne microorganisms"

### Supplementary Figure 3

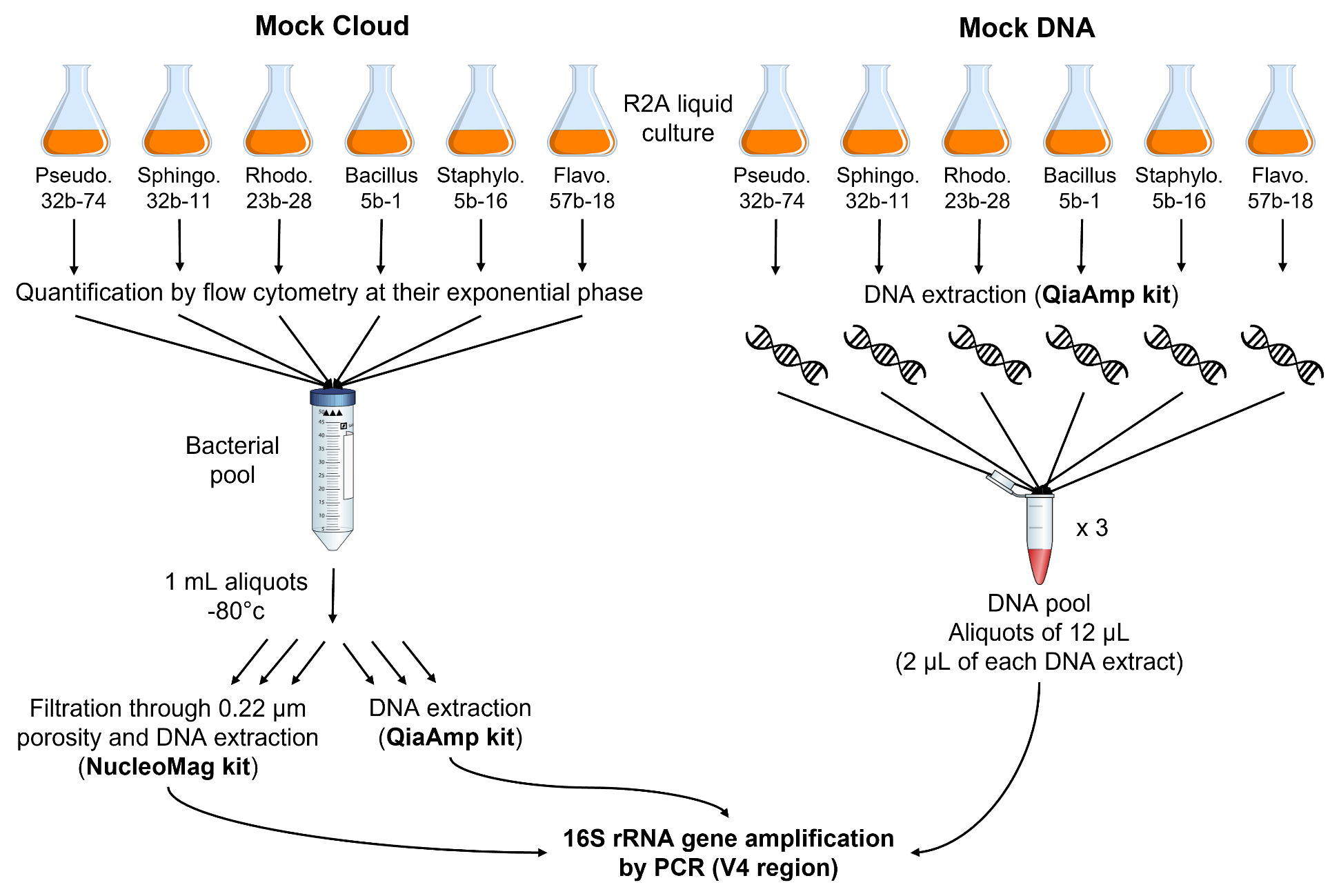

**Supplementary Figure 3:** Protocol for the elaboration of “Mock Cloud” and Mock DNA” samples.
