## Supplementary Figure 4 for "Experimental and methodological framework for the assessment of nucleic acids in airborne microorganisms"

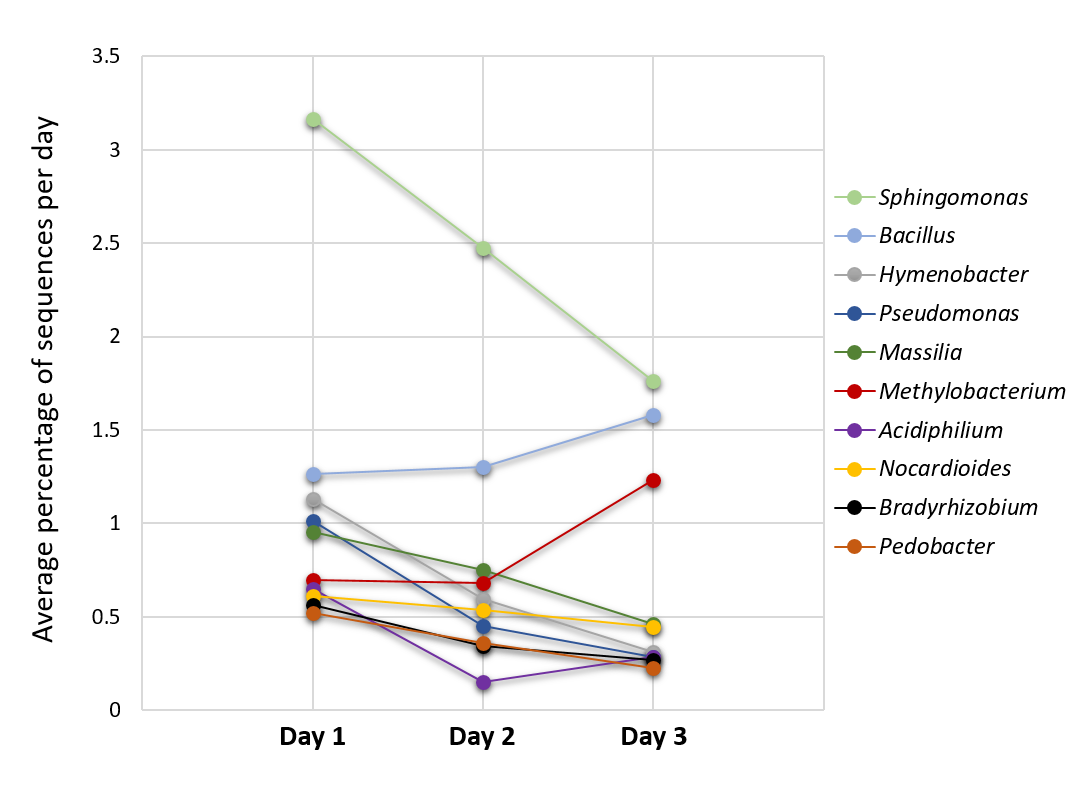

**Supplementary Figure 4:** Average percentages of sequences affiliated with the 10 most abundant genera in aerosol samples.
