## Supplementary Figure 5 for "Experimental and methodological framework for the assessment of nucleic acids in airborne microorganisms"

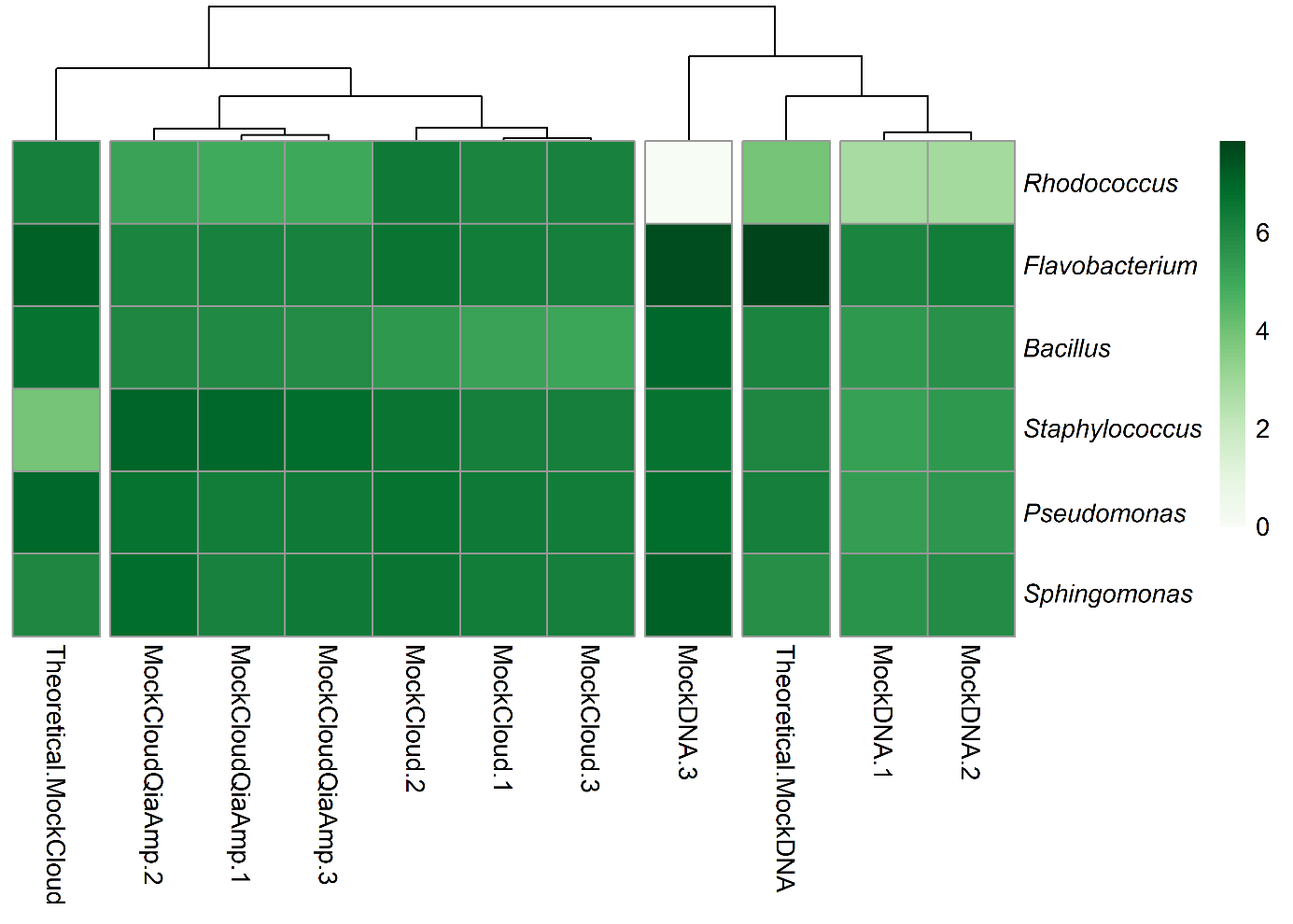

**Supplementary Figure 5:** Distribution of sequence reads in mixed cell suspensions (“MockCloud”) extracted using QIAamp DNA Mini kit or NucleoMag DNA/RNA Water kit (Machery-Nagel), and from mixed DNA solutions (“MockDNA”), and their respective expected (“Theroretical”) distributions based on cell counts and DNA quantifications.
