## Supplementary Table 1 for "Experimental and methodological framework for the assessment of nucleic acids in airborne microorganisms"

**Supplementary Table 1:** Cell and DNA concentrations in the mock communities.

| **Bacterial strain identifier** |  | **Cell concentration in « Mock cloud » sample (cell mL^-1^)** |  | **DNA concentration in « Mock DNA » sample (ng µL^-1^)** |
| --- | --- | --- | --- | --- |
| *Pseudomonas syringae* PDD-32b-74 |  | 1.05×10^8^ |  | 0.25 |
| *Sphingomonas aerolata* PDD-32b-11 |  | 1.05×10^8^ |  | 0.39 |
| *Rhodococcus enclensis* PDD-23b-28 |  | 6.61×10^7^ |  | 0.03 |
| *Bacillus* sp. PDD-5b-1 |  | 4.97×10^7^ |  | 0.13 |
| *Staphylococcus equorum* PDD-5b-16 |  | 3.45×10^6^ |  | 0.14 |
| *Flavobacterium tructae* PDD-57b-18 |  | 1.05×10^8^ |  | 1.01 |
| **TOTAL** |  | **4.35×10^8^** |  | **1.95** |
