## Supplementary Table 2 for "Experimental and methodological framework for the assessment of nucleic acids in airborne microorganisms"

**Supplementary Table 2:** Number of sequencing reads in the samples along data processing.

| **ID sample** | **Raw reads** | **After QC and filters** | **After rarefaction** |
| --- | --- | --- | --- |
| 20200707(1) [Day1.1] | 108 467 | 18 823 | 16 250 |
| 20200707(2) [Day1.2] | 105 748 | 16 250 | 16 250 |
| 20200707(3) [Day1.3] | 126 704 | 24 856 | 16 250 |
| 20200708(1) [Day2.1] | 87 842 | 27 670 | 16 250 |
| 20200708(2) [Day2.2] | 61 384 | 19 027 | 16 250 |
| 20200708(3) [Day2.3] | 89 546 | 24 406 | 16 250 |
| 20200709(1) [Day3.1] | 73 611 | 29 709 | 16 250 |
| 20200709(2) [Day3.2] | 68 747 | 24 392 | 16 250 |
| 20200709(3) [Day3.3] | 72 202 | 23 014 | 16 250 |
| SamplingBlank(1) | 3 916 | 1 771 | - |
| SamplingBlank(2) | 3 564 | 1 870 | - |
| WaterBlank(1) | 3 803 | 2 771 | - |
| WaterBlank(2) | 52 822 | 27 957 | - |
| WaterBlank(3) | 12 113 | 7 890 | - |
| MockCloud(1) | 46 602 | 38 803 | 28 100 |
| MockCloud(1)QiaAmp. | 47 557 | 37 750 | 28 100 |
| MockCloud(2) | 43 476 | 36 225 | 28 100 |
| MockCloud(2)QiaAmp. | 46 671 | 39 828 | 28 100 |
| MockCloud(3) | 33 630 | 28 108 | 28 100 |
| MockCloud(3)QiaAmp. | 41 247 | 33 242 | 28 100 |
| MockDNA(1) | 56 076 | 44 262 | 28 100 |
| MockDNA(2) | 44 544 | 37 578 | 28 100 |
| MockDNA(3) | 36 535 | 30 578 | 28 100 |
